# An Open-Source 3D-Printed Vacuum Manifold for Automated DNA Isolation Improves Yields in the Opentrons Flex and OT-2

**DOI:** 10.64898/2026.09.16.746974

**Authors:** Jack A.T. Dalton, Caleb Goddard, Jean-Baptiste Lugagne

## Abstract

Laboratory automation is increasingly being adopted in academic research to improve throughput, reproducibility, and integration with computational workflows. A common bottleneck in molecular biology workflows is the isolation of DNA, typically performed with spin-column kits relying on centrifugation or vacuum. Existing automation solutions depend on magnetic separation systems requiring costly additional modules and specialist kits, making them impractical for laboratories that perform large numbers of extractions only occasionally. Here we present an open-source, 3D-printed vacuum manifold compatible with the Opentrons family of liquid handling robots that enables automated spin-column DNA isolation for up to 24 samples per run (96 with four manifolds in series). The manifold can be manufactured for approximately $2 using standard ABS filament and is compatible with widely available spin-columns, requiring no additional specialist reagents. Using this system, hands-on time for 24 plasmid minipreps was reduced by 80% compared to manual centrifugation protocols. The automated workflow demonstrated greater plasmid yields and eliminated failed recoveries, though with greater variability in yield between samples. All design files, labware definitions, and Opentrons protocols are released under the CERN Open Hardware Licence or MIT License. This device lowers the cost barrier to automated DNA purification for academic laboratories worldwide.

## 1 Introduction

Laboratory automation is increasingly used within academic research as the need to generate large datasets and perform high-throughput experiments grows due to the increasing adoption of machine learning within biology and existing data becomes a bottleneck on progress[1–3]. Laboratory automation enables the scaling of experiments to much larger sample numbers than would be possible with manual experimentation. Systems such as the Opentrons Flex and OT-2 liquid handling robots have been widely adopted in academic laboratories due to their low cost, open-source software ecosystem, and high customisability [4–6]. Unlike industrial facilities that repeatedly execute identical workflows, academic laboratories tend to use automation platforms across diverse protocols, and are more likely to attempt experimental or unproven protocols that push the boundaries of these technologies. This makes investment in workflow-specific equipment inefficient.

Plasmid DNA isolation (referred to as a miniprep in small scale molecular biology experiments) is one of the most common and time-consuming repetitive tasks in molecular biology [7]. It is a repetitive process that lends itself to automation and can take many hours when processing a large number of samples by hand. Manual protocols using spin-column kits, which rely on centrifugation or vacuum pressure to separate cellular components, are ubiquitous. However, existing approaches to automating this process rely on magnetic-bead purification systems [8]. These require the purchase of a dedicated magnetic module and automation-specific consumables, with costs exceeding $2,000. This makes magnetic-bead automation less accessible for laboratories that perform many minipreps only occasionally but already own a liquid handling robot for other protocols. Spin-columns are also used in a variety of other molecular biology workflows, including genomic DNA isolation, RNA isolation, and PCR purification, so a low-cost, open-source solution for spin-column protocol automation has broad utility.

Vacuum manifolds compatible with spin-columns are well established for manual high-throughput nucleic acid isolation (e.g. Qiagen QIAvac 96), but no low-cost, robot-compatible, open-source design exists for common liquid handlers. To our knowledge, no vacuum-based automated miniprep workflow compatible with commercial liquid handling systems has been previously described.

Here we present an approach that can be adapted to any liquid-handling system and makes use of standard miniprep spin-columns readily available in biology laboratories worldwide, without requiring special equipment beyond a simple 3D-printed device and a vacuum pump or source. By enabling the use of standard, widely available spin-columns on the Opentrons Flex or OT-2, it removes the need for any additional specialised reagents or modules beyond access to a lab vacuum line or commodity vacuum pump. The design is fully open-source and can be reproduced for approximately $2. We show that isolation of plasmid DNA using this system reduces hands-on time by 80%, freeing scientists to focus on other work, while increasing the yield of DNA recovered from the cultures. The manifold can also be used outside the Opentrons on a benchtop vacuum line or in other robotic systems with compatible deck footprints, and the CAD files can be modified to suit alternative platforms.

## 2 Materials and Methods

### 2.1 Design requirements and manifold description

The vacuum manifold was designed against the following requirements. The footprint (127 mm

*×* 85 mm) was chosen to match the Opentrons deck slot. Uniform distribution of wells was required to be compatible with the Opentrons custom labware designer, enabling the creation of custom labware `.json` files that could be integrated into the Python code of the protocol. The manifold was required to be compatible with standard spin-columns with an external diameter of 4 mm at the base. To form a tight vacuum seal with the spin-columns, the manifold was designed with 3 mm pilot holes that were drilled out after printing with a 4 mm drill bit, due to unavoidable deformation during printing. To integrate with 96-well plate protocols, 24 columns per manifold was chosen: this was the largest divisor of 96 that could fit within the footprint of an Opentrons deck slot with sufficient clearance between spin-columns. The manifold was also required to withstand up to 20 mbar of vacuum pressure without deforming.

The resulting device is a 3D-printed manifold that seats up to 24 spin-columns simultaneously within a single Opentrons deck slot. A central barb connects via flexible tubing to an external vacuum source, pulling buffers through the spin-column membranes while the Opentrons pipettes all liquid-transfer steps automatically (Figure 1a). Internal support ribs prevent deformation of the manifold under vacuum pressure (Figure 1b). If the ports do not form a sufficiently tight seal with the spin-columns, electrical tape applied to the top surface and pierced at each column position creates a reliable seal without adhesives. The workflow can be set up with either a 1–50 µL or 50–1000 µL pipette, with the latter providing faster run times. If using less than the full capacity of the manifold, unused column positions are sealed with plugs or tape to maintain vacuum integrity. An overview of the workflow is shown in Figure 1c

**Figure 1.**
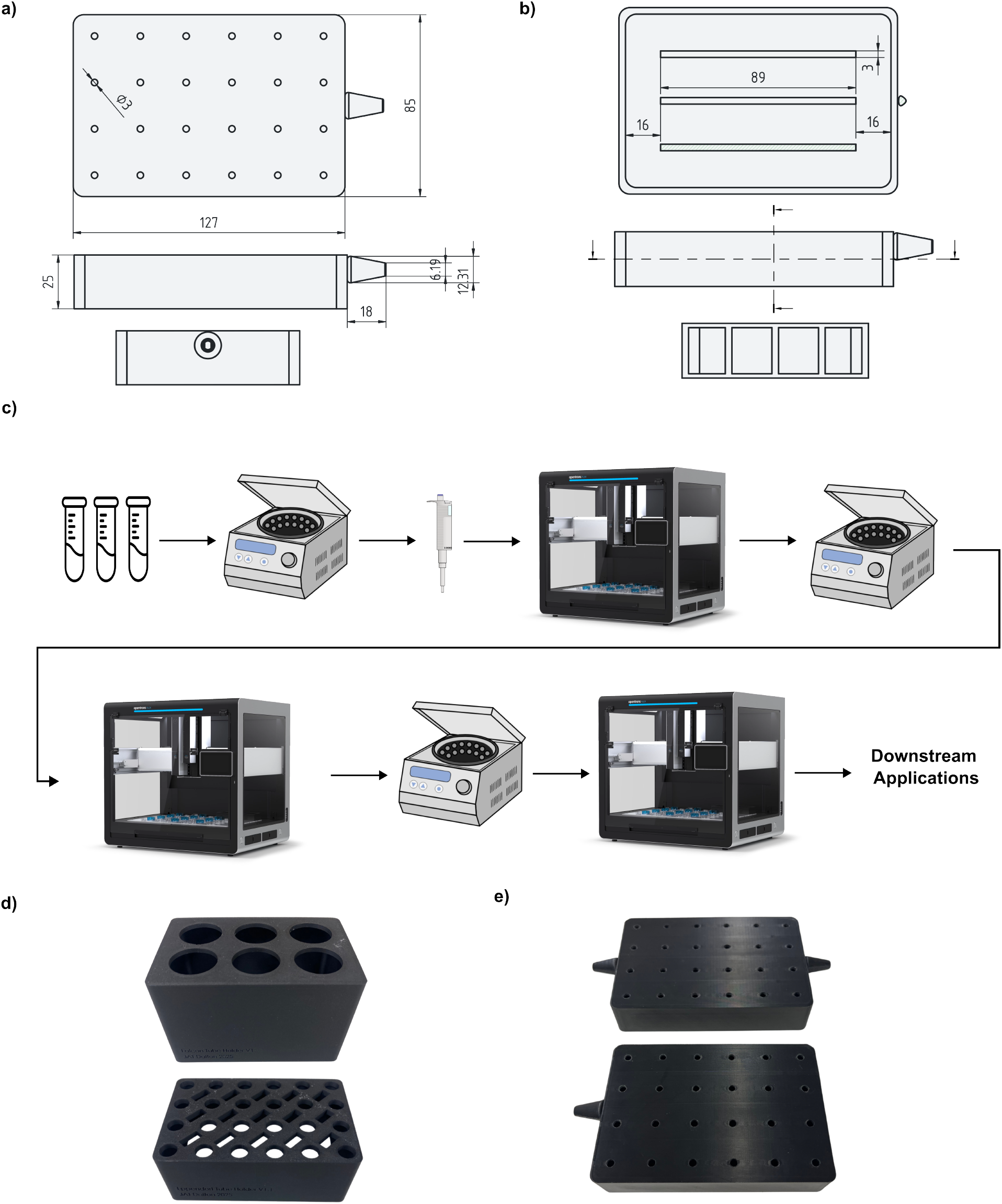
a) Technical drawings of the Opentrons vacuum manifold with key dimensions. b) Cut-through views showing the support ribs added to prevent deformation under vacuum pressure.c) Protocol overview. Slack alerts notify the user when intervention is required. Integration of robot arms with the Opentrons to move samples between different pieces of equipment could eliminate this need. d) The 3D printed Falcon tube and Eppendorf tube racks. e) The final single-port and dual-port vacuum manifolds.

Companion 3D-printed racks for 1.5 mL and 50 mL tubes hold the elution tubes and bulk reagents respectively (Figure 1d). The dual-port version of the manifold features a second barb that enables up to four manifolds to be operated in series within a single Opentrons run, allowing up to 96 samples to be processed (Figure 1e).

### 2.2 Fabrication

The vacuum manifold, 50 mL tube racks and 1.5 mL tube racks were designed in SolidWorks and converted to open formats using FreeCAD 1.1.1. All parts were printed on a Bambu Labs P1S 3D printer with a 0.4 mm nozzle. The tube racks were printed from PLA at 15–20% infill. The vacuum manifold was printed from ABS with 100% infill and four wall loops to prevent gas permeation and increase strength; print time is approximately 3 hours. ABS requires an enclosed printer with good bed adhesion.

The manifold was printed standing on edge, resting on one of the two long side faces (127 *×* 25 mm) that do not carry a barb fitting, rather than lying flat on its largest face (Figure 2a). In this orientation the spin-column face is vertical with the holes pointing sideways, which orients the layer lines around the spin-column mounting holes for greater strength. Printing the manifold flat with the top face down generates support material inside the cavity and produces a weakened surface around each hole from the circular filament paths.

**Figure 2.**
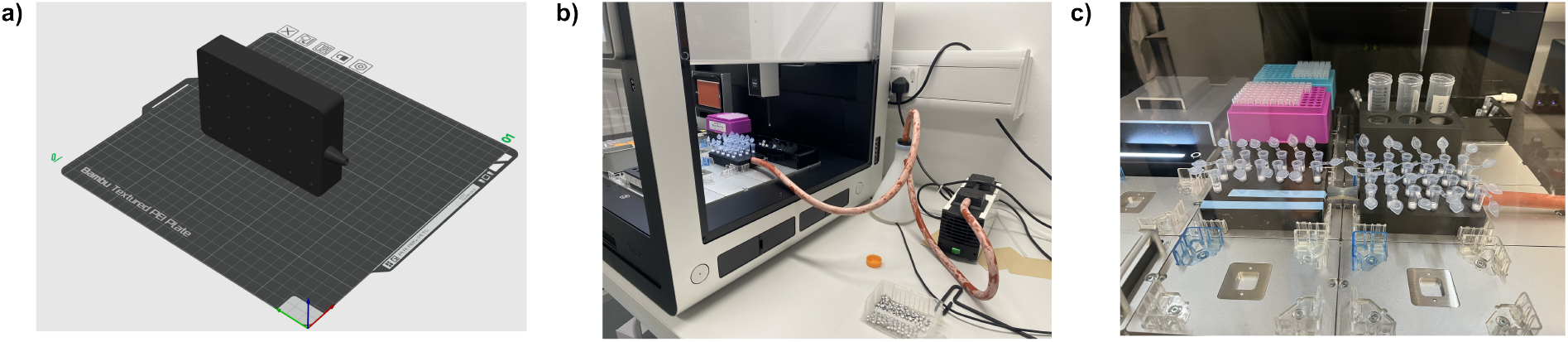
a) View of preparing the manifold for printing in the Bambu Labs slicer. The vacuum manifold should be oriented with the longest side facing down and the largest face oriented sideways as indicated when printing. b) The side panel of the Opentrons must be removed to enable the connection of the vacuum pump to the manifold. This can be done using the hex tool provided with the Opentrons. c) Multiple manifolds can be connected in series to enable the processing of up to 96 samples. Here 37 samples are being processed using two manifolds in series. Tape can be used to seal unused column positions to maintain vacuum integrity.

After printing, each 3 mm pilot hole was enlarged with a 4 mm drill bit and finished with a file, drilling slowly to avoid cracking the ABS. The finished 4 mm holes provide a press-fit seal with standard GeneJET spin-columns. Where the press fit was not sufficient to hold vacuum pressure, strips of electrical self-sealing tape were applied across each row of holes and pierced over each hole using the end of a 1000 µL pipette tip, forming a tight ring of tape around each column position. Flexible tubing was push-fitted onto the manifold barb; the connection is friction-fit and requires no clamp at pressures at or above 20 mbar. Full step-by-step build instructions, slicer settings and the bill of materials are given in Supplementary Information.

### 2.3 Opentrons deck configuration and automated protocol

The automated plasmid miniprep protocol was performed using the Opentrons Flex liquid handling robot, and is implemented in `Miniprep_var_sample_p1000.py` using the Opentrons Python API v2. The custom labware definition `jdvacuum_24_tuberack_500ul.json` was uploaded to the Opentrons App to allow the robot to recognise the manifold well positions during calibration. The deck was configured with a rack for resuspended cell suspensions (slot B2), the vacuum manifold for DNA binding and washing steps (B3), 50 mL Falcon tubes containing lysis, neutralisation, wash and elution buffers (C2), a rack for elution tubes (C3), and tip boxes in positions appropriate to the pipettes in use. Approximately 7 mL of lysis buffer, 9 mL of neutralisation buffer, 25 mL of wash buffer and 2 mL of elution buffer were loaded prior to each 24-sample run. Connecting the vacuum pump to the manifold requires removal of one side panel of the Opentrons using the hex tool supplied with the robot, so that tubing can be routed to the external pump (Figure 2b); alternatively a hole can be drilled through the panel for permanent routing. Overnight cultures (2 mL each) were grown in slanted 14 mL snap-cap tubes at 37°C with shaking at 250 rpm. Cells were transferred to 2 mL Eppendorf tubes and pelleted by centrifugation at 6,800 *×g* for 2 min. All reagents in the following steps were obtained from the GeneJET™ Plasmid Miniprep Kit (Thermo Fisher #K0503). Cells were resuspended by hand in 250 µL Resuspension Solution supplemented with RNase A using a 1000 µL pipette, and the tubes placed in the B2 rack.

The robot then performed lysis and neutralisation in groups of four samples to prevent cells being exposed to lysis buffer for too long; this group size is configurable in the protocol file. Following neutralisation the protocol pauses automatically and notifies the user via Slack. Samples were centrifuged at 21,000 *×g* for 5 min to clarify the lysate, and the supernatant transferred into the spin-columns mounted on the manifold by pouring or pipetting. Vacuum filtration was initiated using a KNF Laboport N 816.3 KT.18 vacuum pump with a catcher flask in the vacuum line, and the protocol resumed to complete the automated wash steps. After washing, the protocol pauses and notifies the user a second time; spin-columns were centrifuged at 21,000 *×g* for 1 min to remove residual ethanol, then transferred to the elution rack (C3). The robot resumed to dispense elution buffer, and samples were centrifuged at 21,000 *×g* for 1 min to recover plasmid DNA.

### 2.4 Manual comparison protocol and yield quantification

The automated protocol was benchmarked against the standard centrifugation-based manual protocol detailed in the GeneJET™ Plasmid Miniprep Kit User Guide (Pub. No. MAN0012655 B), across three independent rounds. Plasmids from the CIDAR and CIDAR Extension MoClo kits, as well as custom plasmids with designed gene fragments inserted into CIDAR backbones [9], were tested.

Eluted plasmid yields were determined using a nanophotometer (Implen NP80) and samples were stored at *−*20°C until further use. A failed recovery was defined as a final concentration below 25 ng/µL. Variability between samples was quantified as the coefficient of variation of yield.

### 2.5 Design files, protocols and data availability

All design files are available at the associated Zenodo repository. STL files are additionally available on Thingiverse to increase discovery. STEP and FreeCAD Python (.py) files are provided as editable and parametric source files, so the design can be regenerated and modified by editing dimensioned parameters (plate size, rib layout, vacuum hole grid, barb geometry) at the top of the build script. STL exports are provided to facilitate direct 3D printing without requiring CAD software. Opentrons custom labware definitions are provided as JSON files compatible with the Opentrons App and Python API v2, and the miniprep protocol is provided as a Python script. A full listing of design files and their licences is given in Supplementary Table S1.

## 3 Results

### 3.1 Iterative design resolved sealing and deformation failures

Initial testing of the automated plasmid miniprep workflow revealed several mechanical and liquid-handling challenges that required redesign of the hardware and protocol steps. The first manifold design lacked adequate sealing and failed to pull lysate through the spin-columns consistently, functioning only when all but one of the outlet holes were sealed, indicating uneven flow resistance across the ports. Applying silicone coatings, and later self-sealing tape, to the ports improved sealing performance. However, the first design also lacked any support within the internal volume, and the resulting pressure difference caused deformation of the manifold top surface under vacuum.

A redesigned manifold (Version 1.2) incorporating internal support ribs (Figure 1b) achieved robust sealing with tape alone, eliminating the need for silicone and preventing deformation. This version provided consistent pull-through across all 24 wells, withstands repeated vacuum cycles at 20 mbar without deformation, and is the design made available in the repository. It is now used routinely in our laboratory to process large numbers of miniprep samples.

### 3.2 Automation reduced hands-on time by 80%

The automated protocol reduced hands-on time from over 2 hours to approximately 40 minutes for 24 samples, an 80% reduction. Total run time was approximately 40 minutes when using the Opentrons 50–1000 µL pipette. Centrifugation steps cannot be automated using the Opentrons alone and require manual intervention at the two protocol pause points and the final elution spin.

### 3.3 Automated isolation gave comparable or greater yields with no failed recoveries

The automated protocol demonstrated comparable or higher plasmid yields at all concentration thresholds tested (25–150 ng/µL) with no failed recoveries (Figure 3a). The manual protocol resulted in a greater percentage of failed recoveries. The Opentrons protocol produced greater or comparable yields of recovered plasmid for all but two plasmids tested (c1j and t14m), with the majority of paired data points falling below the line of equivalence in a pairwise scatter plot (Figure 3b), indicating that for most plasmids the automated method returned higher yields (Figure 3c).

**Figure 3.**
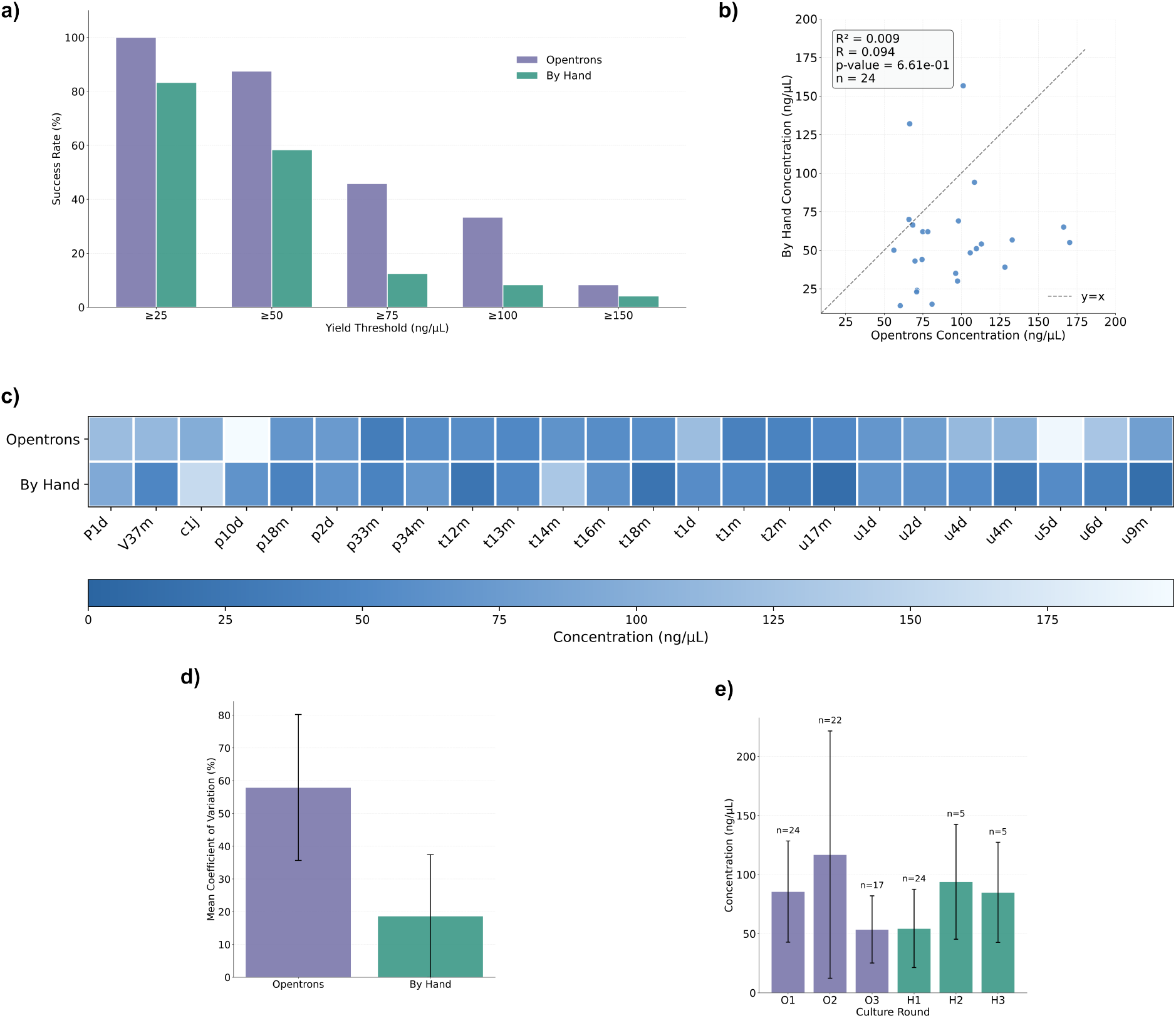
a) Percentage of successful minipreps at varying yield thresholds from 25 to 150 ng/µL. The automated protocol outperforms the by-hand method at all thresholds. b) Scatter plot showing the correlation between yield for the two methods; the majority of data points fall below the line of equivalence, indicating that for most plasmids the automated method resulted in greater yields. c) Heat map comparing individual plasmid yields. d) Bar plot of the mean coefficient of variation; the automated method has a greater coefficient of variation than the by-hand protocol. e) Comparison of different rounds of cultures. Variability changes considerably between rounds for both methods but is elevated in the second automated round.

However, the automated protocol exhibited greater variability in yield than the manual method across the entire library. The mean coefficient of variation was 58% for the automated protocol compared to 19% for the manual protocol (Figure 3d). Most of this variation was observed in the second round of cultures processed on the Opentrons (Figure 3e), whereas the other two automated rounds showed variation in yields comparable to the manual experiments.

## 4 Discussion

The development of an open-source, 3D-printed vacuum manifold for the Opentrons Flex robot effectively addresses the cost barrier associated with automated DNA isolation, particularly for academic laboratories. By utilising readily available spin-column miniprep kits, this low-cost, 3D-printable solution provides a viable alternative to expensive magnetic bead-based systems.

The vacuum manifold can be produced at very low cost. ABS filament costs approximately

$15–20/kg, resulting in a per-manifold print cost of $1–2. All other costs are equivalent to a manual miniprep, as the same reagents are used in the same volumes. By contrast, adopting a magnetic bead-based purification approach requires purchasing an Opentrons magnetic block ($1,750) and a compatible purification kit such as the Zyppy-96 Plasmid MagBead Kit ($400), for a total entry cost exceeding $2,000, along with ongoing specialist reagent costs.

The automated protocol significantly reduced hands-on time for the user by 80%, from over two hours to just 40 minutes for 24 samples, and the total protocol time was optimised to under 40 minutes. Furthermore, the system demonstrated a higher rate of successful plasmid recovery and comparable or greater yields than the standard by-hand centrifugation method, with no failed recoveries observed. While an increase in the coefficient of variation was noted, the higher average yield validates the manifold’s utility.

The principal limitation of the current workflow is that centrifugation steps — cell pelleting, lysate clarification, residual ethanol removal, and final DNA elution — cannot be performed by the Opentrons alone and require manual intervention at protocol pause points. Slack notifications reduce the cost of these pauses by removing the need to monitor the run, but they remain a barrier to fully unattended operation. Integration with robotic arms capable of individual sample manipulation would eliminate these pauses entirely.

## 5 Conclusion

We have presented an open-source, 3D-printed vacuum manifold that enables automated spincolumn DNA isolation on the Opentrons Flex and OT-2 for approximately $2 in materials, using consumables laboratories already own. The device reduces hands-on time for 24 plasmid minipreps by 80% while delivering comparable or greater yields than manual centrifugation and eliminating failed recoveries. This makes automation of DNA purification accessible to a wider community. We hope that the open-source nature of the design encourages the creation and sharing of custom hardware, which is vital for adapting automation systems to novel and experimental academic protocols.

## Data and code availability

All design files, Opentrons labware definitions and protocol code are available at https://doi.org/10.5281/zenodo.21674775 under the CERN Open Hardware Licence Version 2

— Permissive (CERN-OHL-P v2) for hardware and the MIT License for software. STL files are additionally available on Thingiverse.

## CRediT author statement

**Jack A.T. Dalton**: Conceptualisation, Methodology, Software, Design, Investigation, Formal analysis, Visualization, Writing. **Caleb Goddard**: 3D Printing and CAD guidance. **Jean-Baptiste Lugagne**: Conceptualisation, Supervision, Review & Editing.

**Acknowledgements**

J.A.T.D. was supported by the Engineering and Physical Sciences Research Council UK, Centre for Doctoral Training in Engineering Biology, under grant EP/Y034791/1, and by the World Universities Ramsay Postgraduate Scholarship.

## Ethics statement

This work did not involve human subjects, animal experiments, or clinical samples.

## Declaration of generative AI and AI-assisted technologies in the manuscript preparation process

During the preparation of this work, the authors used Claude Code for formatting of LaTeX code as well as the conversion of CAD files to open formats. The authors reviewed and edited the output as needed and take full responsibility for the content of the published article.

## Supplementary Information

### S1 Design files

**Table S1:**
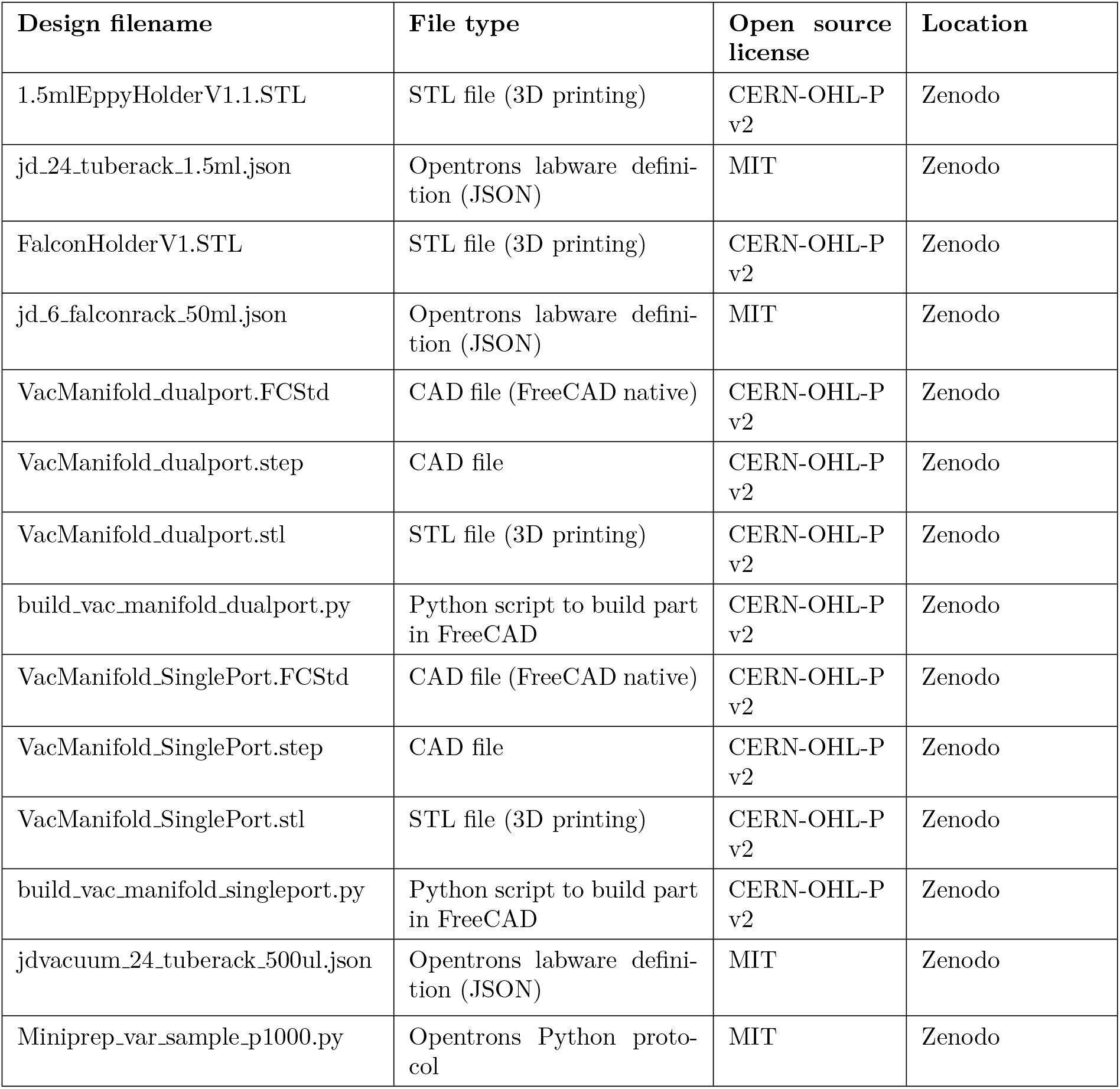
Design files, formats and licences. All files are available at https://doi.org/10.5281/zenodo.21674775.

**VacManifold SinglePort.FCStd / .step / .stl:** Single-port variant of the manifold above, identical geometry apart from omitting the outlet barb, for standalone use or as the last unit in a series

**build vac manifold dualport.py / build vac manifold singleport.py:** FreeCAD Part-API scripts that generate the respective manifolds from dimensioned parameters (plate size, rib layout, vacuum hole grid, barb geometry) rather than fixed geometry, so the design can be re-generated in FreeCAD and modified by editing values at the top of the file.

**1.5mlEppyHolderV1.1.STL:** A 24-well rack for 1.5 mL microcentrifuge tubes sized to fit in an Opentrons deck slot, used to hold the sample source tubes for the miniprep protocol

**FalconHolderV1.STL:** A 6-well rack for 50 mL Falcon tubes sized to fit in an Opentrons deck slot, used to hold the miniprep kit reagents

**jd 24 tuberack 1.5ml.json / jd 6 falconrack 50ml.json:** Opentrons labware definitions for the 1.5 mL tube rack and the 50 mL Falcon rack respectively

**jdvacuum 24 tuberack 500ul.json:** Opentrons labware definition describing the manifold. This definition can be used for both the dual- and single-port versions

**Miniprep var sample p1000.py:** Opentrons Flex protocol for a variable-count (1–24 sample) plasmid miniprep using the vacuum manifold. A P1000 pipette handles lysis, neutralisation, and wash steps while a P50 handles elution, with an optional Slack notification hook. The script requires the requests package, which is pre-installed on the Opentrons Flex; if it is not present in the Python environment the protocol will fail analysis in the Opentrons App. In that case the Slack notification code can be commented out, or the package installed by connecting to the robot over SSH

### S2 Bill of materials

**Table S2:**
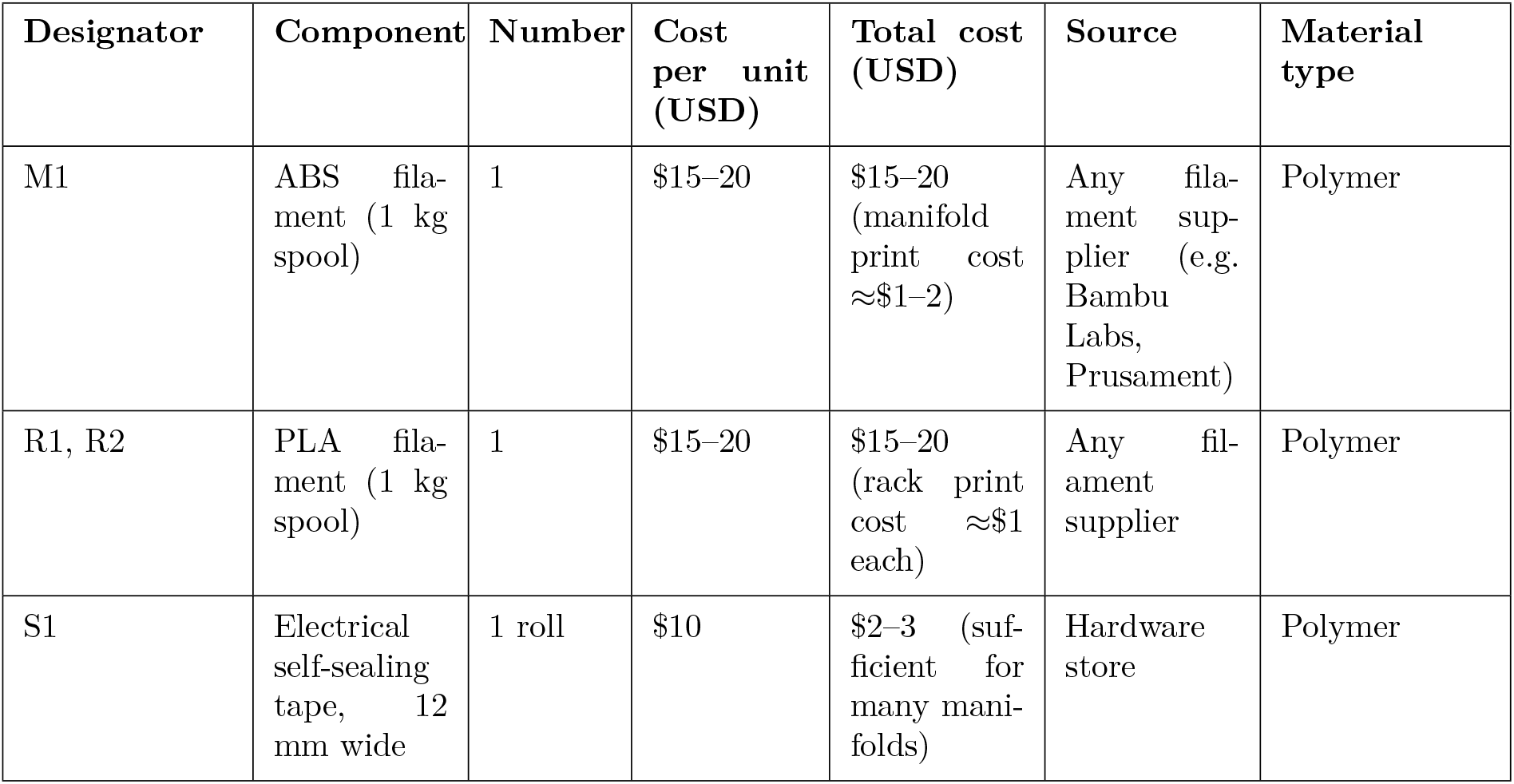
Bill of materials. Reagents (GeneJET Miniprep Kit, standard laboratory consumables) are not listed as they are identical to those used in a manual miniprep. The vacuum pump is assumed to be existing laboratory equipment; any pump capable of reaching 20 mbar is suitable.

### S3 Build instructions

#### Required equipment: Fused filament fabrication 3D printer with ≤0.4 mm nozzle; 4 mm drill bit and hand drill or drill press; file. Alternatively, the manifold and racks can be printed by an online 3D printing service (e.g. Protolabs, Shapeways)

#### Step 1 — Download design files

Download the STL and labware JSON files from the Zenodo repository.

#### Step 2 — Print the vacuum manifold (M1)

Slice VacManifold SinglePort.STL with the following settings:

- Material: ABS (required for solvent resistance and stability under vacuum)
- Nozzle: 0.4 mm
- Infill: 100% (required to prevent gas permeation)
- Wall count: ≥4 perimeters
- Layer height: 0.2 mm (standard)
- Print orientation: standing on edge, resting on one of the long side faces (127 *×* 25 mm) that do not carry a barb fitting. *Do not* print lying flat on the spin-column mounting face.

Print time is approximately 3 hours on a Bambu Labs P1S or equivalent.

#### Step 3 — Print tube racks (R1, R2)

Slice 1.5mlEppyHolderV1.1.STL and FalconHolderV1.STL using PLA, 0.4 mm nozzle, 15–20% infill and the default wall count.

#### Step 4 — Post-process the manifold (pilot holes)

The pilot holes in the manifold top surface are printed at 3 mm diameter to avoid warping. After printing, enlarge each hole with a 4 mm drill bit and finish with a file. Drill slowly to avoid cracking the ABS. The finished 4 mm holes provide a press-fit seal with standard GeneJET spin-columns (4 mm external base diameter).

#### Step 5 — Apply electrical tape (S1)

If the press fit of the spin-column is not tight enough to hold vacuum pressure, cut strips of electrical tape to span each row of holes on the manifold top surface and press firmly. Using the end of a 1000 µL pipette tip, pierce the tape over each hole. Check each hole with a spare spin-column; the column should seat firmly and not rock.

#### Step 6 — Attach vacuum tubing

Push flexible tubing onto the barb on the side of the manifold. The connection is friction-fit; no clamp is required for pressures at or above 20 mbar. The barb can be redesigned in CAD to accommodate different tubing diameters.

#### Step 7 — Install Opentrons labware definition

Upload jdvacuum 24 tuberack 500ul.json to the Opentrons App via *More* → *Custom Labware*.

**Safety note:** ABS fumes can be irritating; print in a well-ventilated area or an enclosed printer with a HEPA/activated carbon filter. Drilling ABS produces fine particulate; wear appropriate eye protection

### S4 Operation checklist

1. Resuspend cell pellets by hand with 250 µL Resuspension Solution (with RNase A) using a 1000 µL pipette, and place 1.5 mL tubes in the B2 rack.
2. Load the deck as described in Methods and upload Miniprep var sample p1000.py to the Opentrons App.
3. Start the protocol. The robot performs lysis and neutralisation in groups of 4.
4. While the robot is running, place spin-columns into the manifold.
5. *First manual pause (Slack notification):* Centrifuge all samples at 21,000 *×g* for 5 min to clarify the lysate. Transfer the supernatant into the spin-columns on the manifold. Activate the vacuum and resume the protocol.
6. The robot performs the wash buffer steps.
7. *Second manual pause (Slack notification):* Centrifuge spin-columns at 21,000 *×g* for 1 min to remove residual ethanol. Place columns into the corresponding elution tubes and resume. Do not skip this spin step, as residual ethanol will significantly reduce yields.
8. The robot dispenses elution buffer.
9. *Final manual step:* Centrifuge at 21,000 *×g* for 1 min to recover plasmid DNA into the elution tubes.
10. Quantify yield using a nanophotometer. Store at −20°C.

